# Thermally controlled state switches for macrophage immunotherapy

**DOI:** 10.1101/2025.06.09.658763

**Authors:** Ann Liu, Abdullah S. Farooq, Mohamad H. Abedi, Cameron A. B. Smith, Di Wu, Mikhail G. Shapiro

## Abstract

Advances in cellular immunotherapy promise new treatments for conditions such as cancer, autoimmune disease, and heart disease. While engineered cells have the ability to recognize clinically relevant signals, traffic to disease sites and interface with the host immune system, their activity must be tightly controlled to minimize undesirable effects in healthy tissues. One approach to obtaining specificity is to activate the cells spatially using externally applied energy, such as ultrasound-delivered heating. To facilitate such control, we designed and characterized a genetic circuit that enables stable transcriptional activation of macrophages after a brief thermal stimulus, resulting in the expression of reporters or secretion of the cytokine IL-12. We demonstrate that *in vivo* activation of a mouse macrophage cell line containing this bioswitch results in spatially localized gene expression for at least 14 days after ultrasound heating. This thermal bioswitch provides a precise control element for cell-therapeutic agents.

## INTRODUCTION

Cellular immunotherapy has shown tremendous clinical promise in cancer and autoimmune disease, with therapies such as chimeric antigen receptor (CAR)-T cells providing high efficacy in hematologic malignancies^1,2^ and B cell-based autoimmunity^3–5^. However, significant challenges have arisen in generalizing cell-based treatments to additional diseases and cell types. For example, in solid tumors, the microenvironment is highly immunosuppressive, limiting the ability of therapeutic cells to persist and execute their cytotoxic activity^6^. To combat this issue, strategies such as engineering “ armored” cells to express cytokines that promote an inflammatory microenvironment have been explored^7^. However, the secretion of these cytokines in off-target tissue or into systemic circulation can cause widespread inflammation outside of the tumor, resulting in potentially severe side effects^8,9^.

To mitigate the toxic effects of systemically active cell therapies, methods for local control of transgene expression have been developed. While optogenetic approaches have been explored, their *in vivo* use is limited due to the poor tissue penetration of light^10,11^. Recently, our group and others have demonstrated thermal control of CAR-T receptor and cytokine expression in T cells^12–14^. Using non-invasive methods such as ultrasound therapy, thermal energy can be deposited deep inside tissues with a high degree of spatial precision^15^. Localized tissue heating has been used to actuate cytokine release from engineered bacteria and T cells *in vivo*^13,16^.

Here, we extend the toolkit for thermal remote control to macrophages and enable sustained activity. Due to the documented ability of macrophages to infiltrate deep into and persist within solid tumors, this cell type is a promising candidate for future cellular immunotherapies. Although tumor-associated macrophages typically promote tumor growth and immunosuppression^17^, engineered macrophages expressing CARs, pro-inflammatory cytokines or other therapeutic effectors can be reprogrammed as anti-tumor agents^18–20^. To enable thermal remote control of macrophages, we introduce a genetic circuit with low baseline activity and high, sustained activation in response to a brief thermal stimulus. We validate and characterize the performance of this bioswitch in a macrophage cell line (RAW 264.7) and demonstrate precise control over reporter genes and a secreted cytokine. Then, using a clinically approved ultrasound therapy device, we demonstrate sustained *in vivo* activation of the engineered macrophages in a subcutaneous tumor model. This switch extends the toolbox of synthetic biological tools for immune cell engineering to macrophages and provides a robust method for sustained functional activation *in vitro* and *in vivo*.

## RESULTS

### Constructing a temperature-sensitive state switch architecture in macrophages

We constructed a basic genetic circuit consisting of an actuator element that responds to elevated temperature by expressing Cre recombinase and a toggle switch encoding distinct genes in the OFF and ON states. In the actuator element, we placed a heat shock protein (HSP) promoter^12–14,21^ upstream of the recombinase. In the toggle switch, we used a constitutive EF1*α* promoter to drive a flip-excision switch^22^ consisting of a red fluorescent protein (RFP) immediately upstream of an inverted enhanced green fluorescent protein (GFP). When Cre is expressed, the target DNA between the cut sites is irreversibly inverted, permanently switching the circuit from OFF (RFP expression) to ON (GFP expression) (Figure 1a).

**Figure 1.**
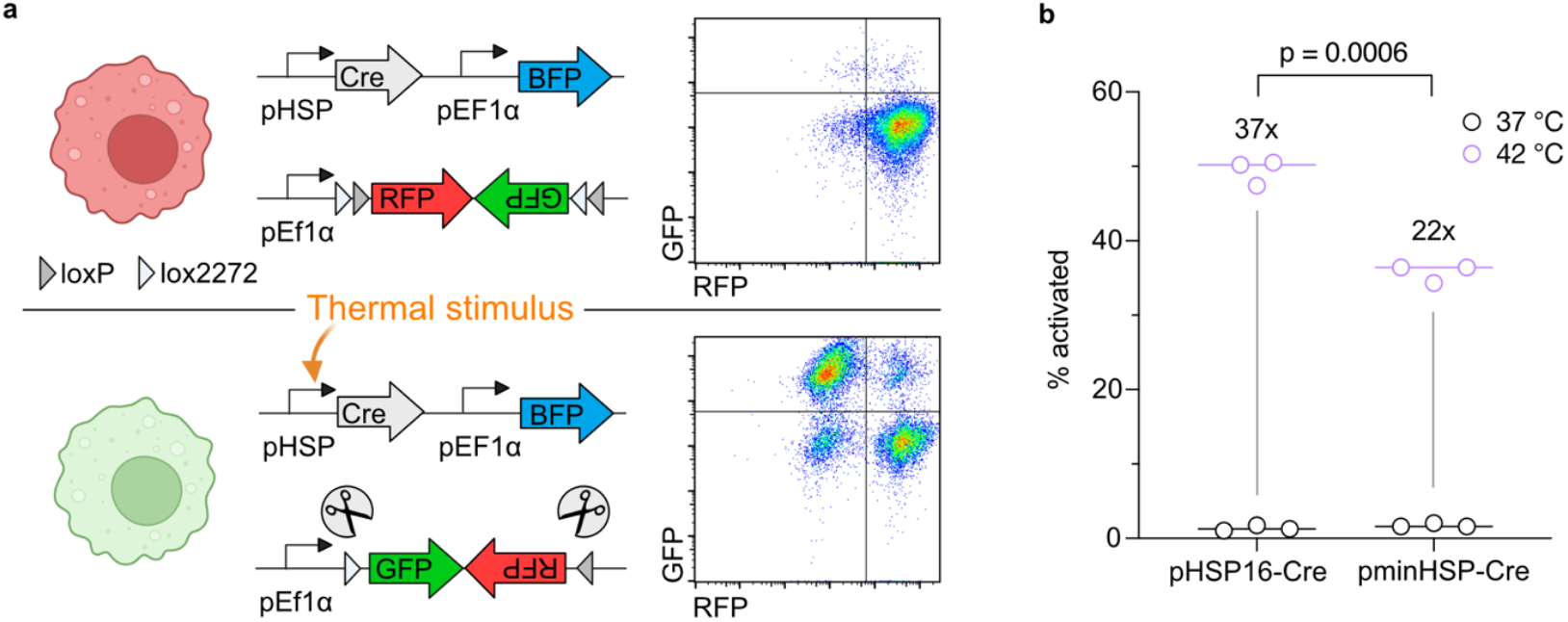
Construction and evaluation of a temperature sensitive state switch in macrophages. (**a**) Diagram illustrating the genetic circuit architecture in the OFF state (top) and ON state (bottom) after a thermal stimulus is applied. Representative flow cytometry plots shown on the right. (**b**) Percent activated cells (eGFP+/BFP+) after a 1 hour incubation at either 37 or 42 °C, measured via flow cytometry. Cells were first gated for BFP+. Fold change between 37 and 42 °C is listed above sample.

We first selected two candidate HSP promoters to compare – the HSP16F promoter identified in previous work^12,23^ and a synthetic minimal HSP promoter (minHSP). The minHSP promoter consists of 4 consecutive repeats of the heat shock element (HSE) 5’-nGAAnnTTCn’-3’ followed by a minimal TATA-box promoter with low basal activity. The pHSP-Cre constructs also included a blue fluorescent protein (BFP) transduction marker.

Each pHSP-Cre construct was lentivirally transduced into RAW 264.7 macrophages along with the fluorescent reporter toggle switch, and the cells were sorted to generate stable polyclonal cell lines. We evaluated the performance of each candidate HSP promoter by incubating the engineered cells at 42 °C for 60 minutes and then measuring the fraction of cells in the ON state 24 hours later using flow cytometry. Both HSP promoters demonstrated low baseline activation in the absence of thermal stimulation, with < 2% of cells in the ON state (Figure 1b). The HSP16F promoter demonstrated higher activation at 49.7%, and we selected it for further experiments.

### Characterizing thermal switch activation in vitro

After finalizing our bioswitch construct, we generated a stable RAW 264.7 cell line containing the HSP16F-Cre and fluorescent reporter toggle switch constructs (RAW-togFP) and proceeded to characterize its response to different thermal induction conditions. We transiently heated the RAW-togFP cells at temperatures ranging from 37 to 43 °C for 1 hour and then used flow cytometry to measure the proportion of cells in the ON state. The switch demonstrated a small amount of activity at 41 °C and a significant jump in activation at 42 °C (Figure 2a). Importantly, RAW-togFP cells exhibited low baseline activity between 37 and 40 °C, which encompasses physiological and febrile body temperatures^24^. We observed that the vast majority of cells were not viable after induction at 43 °C for 1 hour.

**Figure 2.**
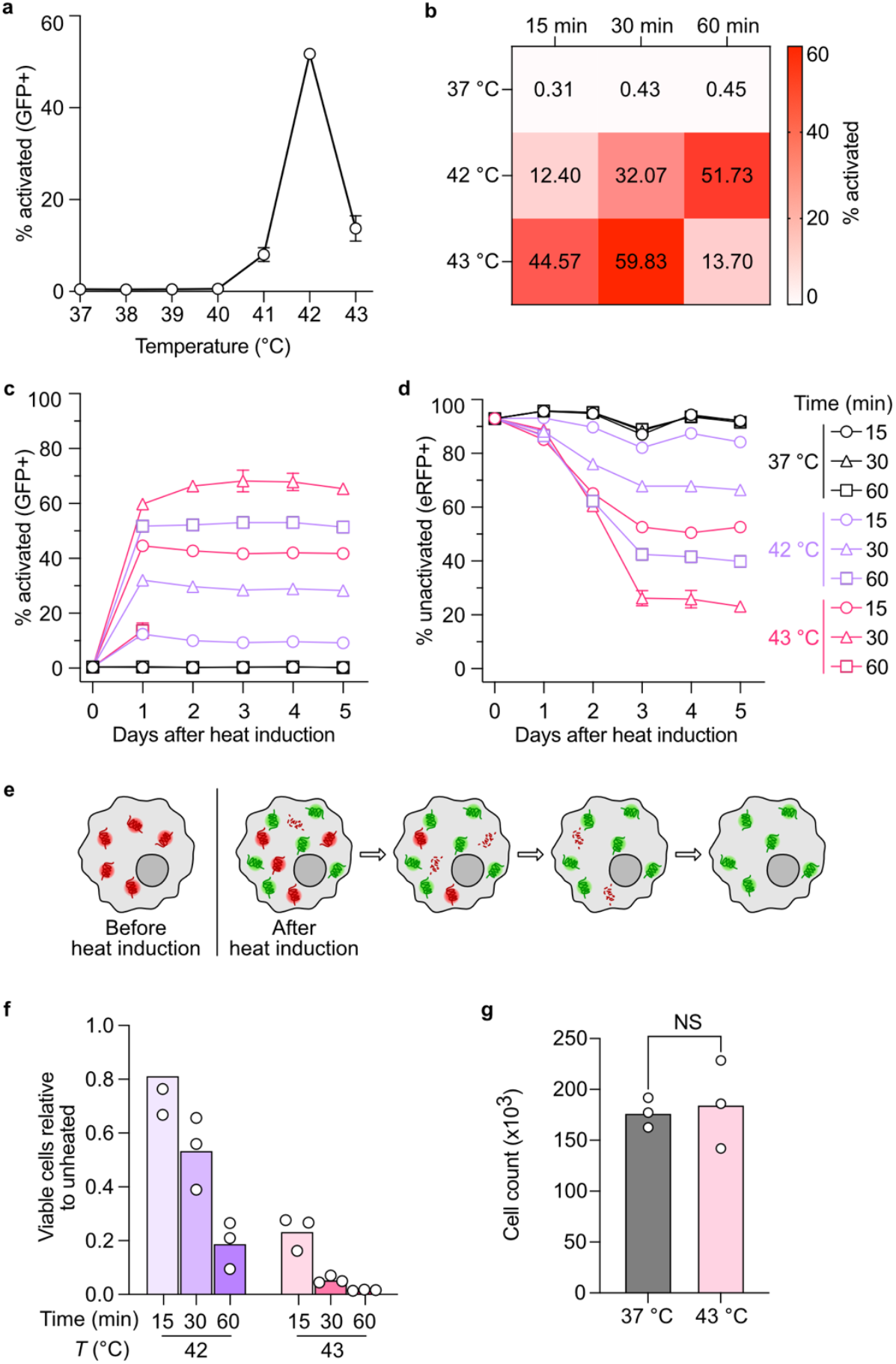
Characterizing induction conditions of the temperature sensitive state switch in macrophages. (**a**,**b**) Percent activation of engineered cells after 1 hour of induction at 37-43 °C (a) and 15-60 minutes of induction at 37, 42, or 43 °C (b). (**c**,**d**) Kinetics of GFP expression (c) and emiRFP670 (eRFP) expression (d) over 5 days following 15-60 minutes of induction at 37, 42, or 43 °C. Cells were first gated for doublet discrimination and BFP+. (**e**) Graphical representation of GFP and eRFP kinetics after heat induction. (**f**) Viable cells relative to unheated cells 24 hours after 15-60 minutes of induction at 42 or 43 °C were counted using flow cytometry after staining with 7-AAD. (**g**) Four days after 15 minutes of induction at 37 or 43 °C, an equal number of cells from either condition were seeded. Viable cells were stained using 7-AAD and counted 24 hours later using flow cytometry. n = 3 for all panels. Bars represent SEM.

To further optimize the heating paradigm, we investigated the effect of varying the duration of heating on switch activity and cell viability. After incubating cells at 42 or 43 °C for 15, 30, or 60 minutes, we observed a positive correlation between induction duration and switch activation (Figure 2b), with the exception of the 43 °C 60 minute condition. To assess the kinetics and stability of circuit activation, we measured the proportion of cells in the ON and OFF states for five days following heat induction. At each condition, activation reached or approached its maximum after 24 hours and remained stable for at least five days (Figure 2c). Cells incubated at 37 °C remained at well below 1% activation for all induction durations. The expression of the OFF state marker (eRFP) decreased steadily until three days post-induction, and then persisted at a stable level through day five (Figure 2d). This corresponds to the gradual degradation of pre-expressed eRFP proteins, as the intracellular half-life of fluorescent proteins is 20-30 hours^25^ (Figure 2e). Five days after induction, almost all GFP+ activated cells were also eRFP−, indicating that every copy of the switch construct in a given cell is inverted by the Cre recombinase (Figure S1). Thus, the activation of the thermal circuit from the OFF to the ON state appears to be a high-performance state switch, with minimal baseline leak at physiological temperatures and sustained activity over five days after activation. We also observed that the transient heat stress did not cause loss of the genetic circuit itself, as evidenced by the stable expression of the BFP marker in heated and unheated RAW-togFP cells (Figure S2).

In addition to characterizing circuit activity and kinetics, we sought to investigate the effect of transient heat induction on RAW 264.7 cell viability. While 42 and 43 °C are well below the temperatures used for thermal ablation^26^, there may still be adverse effects on cell health^27^. As expected, higher temperatures and longer incubation durations both resulted in decreased cell viability 24 hours after heat induction (Figure 2f). To facilitate *in vivo* experiments, we favored a short induction time and selected 43 °C for 15 min as the induction condition. To ensure that the immediate toxicity of the heat induction does not have a lasting effect on the proliferating cell population, we compared the growth rate of cells incubated at 37 or 43°C for 15 min and found that the growth rate recovered to comparable levels by day 4 after heating (Figure 2g).

### Sustained activation of thermal switch in vivo via ultrasound-induced hyperthermia

To demonstrate external thermal control of engineered RAW 264.7 cells *in vivo*, we designed a bioluminescent version of the genetic circuit, togLuc (Figure 3a). We replaced BFP with Antares^28^, a fusion of the luciferase NanoLuc and the orange fluorescent protein cyOFP, and placed the luciferase AkaLuc^29^ in the inverted portion of the switch construct with GFP. This system uses Antares to image all engineered cells and ON-state AkaLuc to monitor switch activation^30^. The inclusion of GFP enabled *ex vivo* analysis using flow cytometry and fluorescence microscopy.

**Figure 3.**
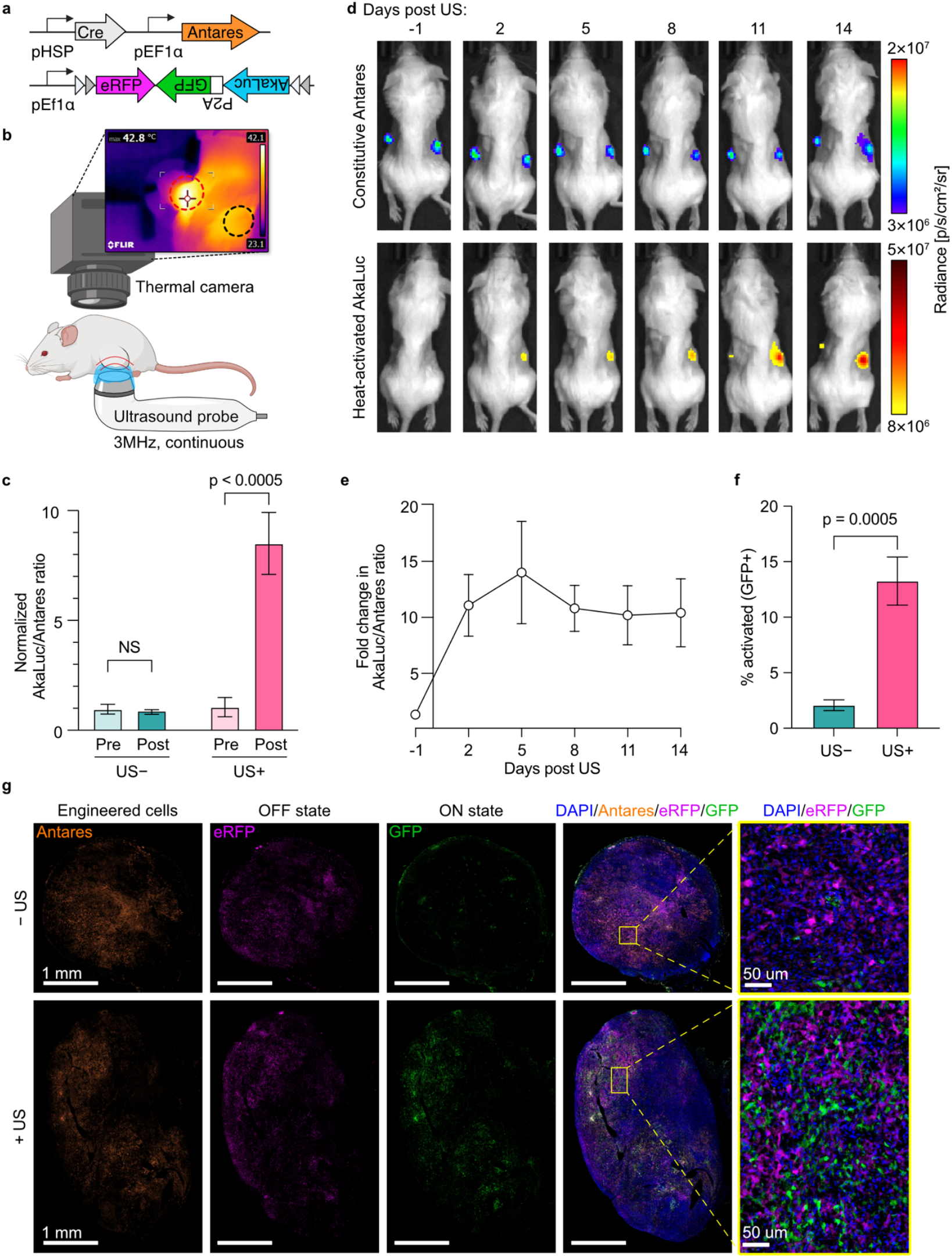
Ultrasound-induced hyperthermia activates engineered macrophages in vivo with long-term persistence. (**a**) Genetic circuit architecture of luminescent reporter thermal switch. (**b**) Graphic of ultrasound treatment set up of mouse with bilateral flank tumors. Thermal image shows mouse during US of left flank tumor (red circle = treated tumor, black circle = untreated tumor). (**c**) Activation of RAW cells bearing thermal switch one day before (Pre) and two days after (Post) US treatment in vivo. Circuit activation was quantified by the AkaLuc/Antares bioluminescence ratio, and normalized to the “ Pre, US−” condition. N = 12 mice. (**d**) Luminescence images of one representative mouse bearing bilateral subcutaneous tumors containing engineered RAW cells, with right-sided tumor treated with US on day 0. Top row: signal from constitutively expressed Antares activity. Bottom row: signal from AkaLuc activity. (**e**) Induction of AkaLuc expression in the US treated tumor quantified as the AkaLuc/Antares ratio normalized to the untreated contralateral tumor. n = 7 mice. (**f**) Percent of engineered RAW cells flipped ON in the treated (US+) or untreated (US−) tumors 14 days after treatment, as measured by flow cytometry. Cells were first gated for doublet discrimination, dead cell staining, and Antares+. n = 5 mice. (**g**) Fluorescence microscopy of tumor sections collected 14 days after US. Bars represent SEM.

We subcutaneously implanted 4T1 breast tumor cells mixed with RAW-togLuc cells bilaterally in the flanks of Balb/C mice. Once tumors reached ~100 mm^3^, baseline bioluminescence imaging was performed, and thermal activation took place the following day. We randomly assigned either the left- or right-sided tumor to be heated using ultrasound, with the contralateral tumor serving as an unheated control. To heat the tumors, we used a commercially available human ultrasound therapy device emitting a continuous, unfocused ultrasound beam at a frequency of 3 MHz from a 1 cm^2^ circular transducer. The transducer was positioned below the tumor and an infrared thermal camera was used to monitor the skin temperature from above (Figure 3b). The power output from the ultrasound device was manually adjusted between 0.1-2.0 W/cm^2^, an applied acoustic peak-to-peak pressure of 181-516 kPa (Fig S3), to maintain a skin temperature of 43 ± 1 °C for 15 minutes, guided by the live thermal imaging.

Two days after US treatment, the AkaLuc/Antares ratio from US-treated tumors increased by over 8-fold compared to pretreatment, while untreated tumors showed no significant change (Figure 3c). AkaLuc signal, which is switched on by heating, was normalized to the constitutive Antares signal from each tumor to normalize for tumor size, light attenuation, and RAW cell density.

We assessed the stability of the thermally activated state *in vivo* over 14 days following heating (Figure 3d). US-treated tumors maintained significantly higher AkaLuc signal (ON state) compared to US-negative control tumors, with a steady increase over time (Fig S4a). To eliminate potential confounds from increases in tumor size, the AkaLuc/Antares ratio was used to calculate the fold-change in ON signal from treated vs untreated tumors. We observed that the population of thermally activated RAW-togLuc cells remained remarkably stable for at least two weeks (Figure 3e). Notably, treated and untreated tumors exhibited similar growth curves, indicating that US treatment itself did not significantly affect tumor growth (Figure S4b).

After the last imaging session on day 14, we collected tumors for flow cytometry and observed that 13.3 ± 2.2% (N = 5, mean ± SEM (standard error of the mean)) of RAW-togLuc cells in the US-activated tumors were in the ON state, compared to 2.1 ± 0.5% in the untreated tumors (Figure 3f). Tumors from one mouse were fixed for cryosectioning and fluorescence microscopy (Figure 3g). GFP+ cells (indicating activated RAW-togLuc cells) in the treated tumor were evenly distributed throughout Antares+ regions, suggesting that thermal activation occurred uniformly within the tumor. Regions lacking any fluorescence likely correspond to unlabeled 4T1 cells. No cells were found to be both GFP+ and eRFP+ (Figure 3g, right column), verifying that the circuit operates as a binary switch *in vivo*.

### Thermal control of cytokine release from macrophages harboring genetic switch

After characterizing the thermal switch both *in vitro* and *in vivo* using fluorescent and bioluminescent reporters, we adapted this switch to control the expression of a secreted therapeutic output. IL-12 is a potent cytokine that promotes a proinflammatory microenvironment by enhancing T cell responses and inhibiting immunosuppressive cells, but its systemic delivery can cause severe toxicity, including neutropenia, neurotoxicity, and death^31^. Temperature sensitive state switches could enable remote control of IL-12 release in only the desired anatomical locations. To enable this future application of our state switches, we set out to demonstrate this triggered release capability *in vitro*.

We created a new construct placing GFP immediately upstream of an inverted IL-12, which was followed by an internal ribosome entry sequence and eRFP (Figure 4a). We incubated RAW-togIL12 cells at 37 or 43 °C for 15 minutes and measured the amount of IL-12 secreted into the culture media. In a 24 hour period, over 50 ng of IL-12 was produced from 500,000 heated cells, compared to 0.5 ng produced from unheated cells (Figure 4b). To verify that the IL-12 measured from heated RAW-togIL12 cells was specific to activation of the togIL12 circuit rather than endogenous IL-12 production or an effect of lentiviral transduction or heating, we generated a RAW-Ctrl cell line transduced with only the IL-12 toggle switch and lacking the HSP-Cre actuator (Figure S5a). Heated RAW-Ctrl cells produced negligible amounts of IL-12 compared to heated RAW-togIL12 cells (Figure S5b).

**Figure 4.**
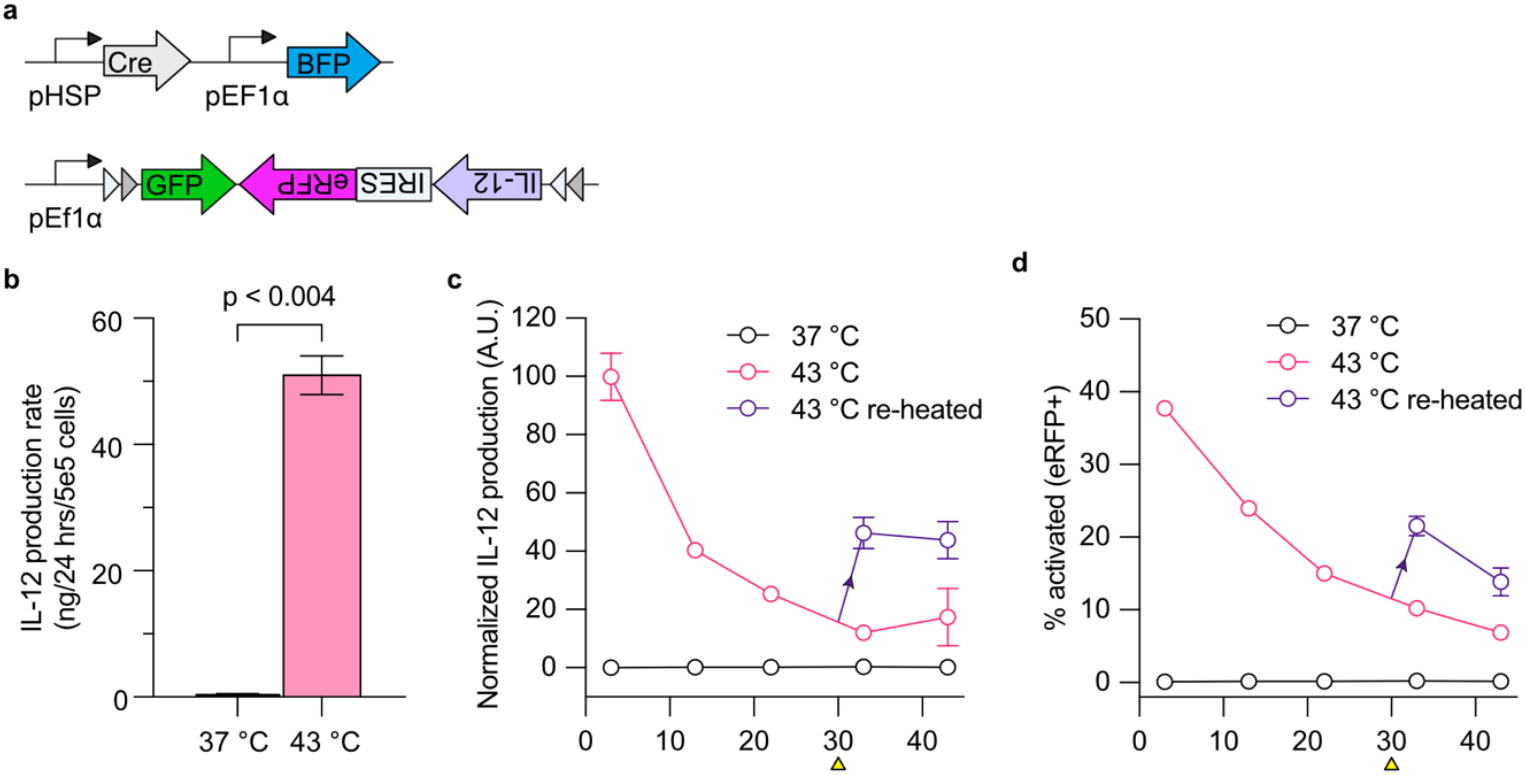
Temperature controlled state switches turn on sustained IL-12 production from engineered macrophages. (**a**) Diagram illustrating togIL12 genetic circuit architecture. (**b**) IL-12 production over 24 hours from RAW-togIL12 cells incubated at 37 or 43 °C for 15 minutes. (**c**,**d**) Kinetics of IL-12 production (c) and percent activation (eRFP+/BFP+) of RAW-togIL12 cells (d) over 43 days following incubation at 37 or 43 °C for 15 minutes at t = 0 days. At t = 30 days (yellow arrowhead), a subpopulation of cells that had been heated at 43 °C at t = 0 were incubated again at 43 °C for 15 minutes and then further assessed on days 33 and 43 (purple lines). (**c**) At each time point, culture media was collected from cells that had been seeded 3 days prior for IL-12 quantification. IL-12 was normalized to cell count from flow cytometry, and then normalized to the peak value across all days. (**d**) For flow cytometric analysis, cells were first gated for doublet discrimination and BFP+. n = 3 for all panels. Bars represent SEM.

We assessed the kinetics of IL-12 expression from RAW-togIL12 cells over 43 days after a single heat induction. IL-12 production remained significantly elevated relative to unheated controls for 22 days, despite a gradual decline over time (Figure 4c). Flow cytometry revealed a similar trend, with the activated cell population peaking at 37.7% before declining (Figure 4d). We reasoned that the continuous expression of the IL-12 transgene may cause activated cells to experience metabolic burden, resulting in progressive epigenetic silencing of the ON state toggle switch element^32^ or selection pressure against activated cells. Unheated and heated RAW-togIL12 cells show similar BFP (HSP-Cre construct transduction marker) and GFP (OFF state toggle switch construct) expression dynamics over the 43 day time period (Figure S6a,b), indicating that OFF state cells maintained both circuit components. A small decline in both fluorophores could be attributed to epigenetic silencing of genomically integrated transgenes^32^.

After observing the decline in IL-12 during the time course, we tested whether an additional round of heating could restore IL-12 production via activation of the remaining OFF state cells. Thirty days after the initial thermal induction, a subpopulation of heated RAW-togIL12 cells were incubated again at 43 °C for 15 minutes. The RAW-togIL12 cells heated only once at t = 0 continued to be assessed in parallel with the re-heated cells. Three days later (t = 33 days), IL-12 produced from the re-heated cells jumped from 12.1 ± 1.9% to 46.3 ± 5.3% (N = 3, mean ± SEM) of the peak value from the original RAW-togIL12 cells (Figure 4c, purple line). The fraction of cells in the ON state also increased from 10.2 ± 0.9% to 21.5 ± 1.3% three days after the second heating (Figure 4d, purple line). From both IL-12 expression kinetics and circuit activation, we observed that a second induction resulted in approximately 50% of the maximum expression and activation from the first induction. This is consistent with the observation that at 30 days, approximately 50% of heated RAW-togIL12 cells were in the OFF state and eligible to be activated (Figure S6b). These results show that repeated heating can augment expressed payload secretion, enabling controlled, repeatable dosing.

## DISCUSSION

The temperature-activated state switch described in this study enables rapid activation of long-term gene expression in macrophages in response to transient temperature elevations. When this switch is used to actuate the release of IL-12, macrophages are able to produce IL-12 for up to three weeks, while a second heat treatment 30 days after the initial induction can switch on a new population of macrophages, enabling the possibility of carefully tuned dosing of this cytokine in a therapeutic context. Our characterization of this switch provides the basis for a new toolkit to enable precise control over gene expression in macrophage immunotherapy models. Furthermore, as a proof of concept, we demonstrate remote control of our state switch in engineered macrophages in vivo using ultrasound-mediated hyperthermia.

Future work will include deploying these IL-12 switches in bone marrow derived macrophages (BMDMs) to more closely resemble cell therapy models, and then demonstrating therapeutic efficiency in a murine tumor model. While this study centers on cancer immunotherapy applications, a noninvasively controlled state switch in macrophages has potential applications in many contexts, such as the study of macrophage biology and control of other therapeutic cell types.

Our study adds to the growing body of work on temperature-controlled cellular immunotherapies^13,14,20^. While effective, these approaches require up to six repeated treatment sessions to maintain expression of the heat-inducible payload, which can be burdensome to both the patient and the hospital system providing treatment. Our permanent switch circuit removes this burden by enabling long-term activity following a single 15-minute session.

One limitation of this stable switch activation is the possibility of activated cells continuing to express the therapeutic payload once treatment is no longer needed, or activated macrophages migrating to other locations in the body. A potential strategy to overcome this is adding a kill-switch to the engineered macrophages that can be activated pharmacologically, such as an inducible caspase system^33^. With these improvements, thermal bioswitches have the potential to help macrophages and other engineered cell types tackle the challenge of action specificity.

## METHODS

### Plasmid construction

All plasmids were designed using SnapGene (GSL Biotech) and constructed using Gibson Assembly using enzymes from New England Biolabs. The minimal HSP promoter was amplified by PCR from pGL4.41[luc2P/HSE/Hygro] (Promega). The IL-12b gene (NCBI Accession Number: NM_001303244.1) was synthesized by IDT. Constructs were transformed into NEB Turbo and NEB Stable E. coli (New England Biolabs) after assembly for plasmid preparation and sequence verified. The fluorescent reporter toggle switch construct for initial screening of HSP promoters contained the red fluorescent protein mScarlet, which was replaced with the far-red fluorescent protein, emiRFP670 (eRFP), for the remainder of the study for logistical reasons.

### Cell lines

4T1 mouse mammary carcinoma cell line was obtained from the American Tissue Culture Collection (ATCC) and cultured in RPMI 1640 media (ATCC) supplemented with Penicillin/Streptomycin (Corning) and 10% FBS (Bio-Techne). RAW 264.7 mouse macrophages were obtained from ATCC and cultured in DMEM media (ThermoFisher) supplemented with 1X Penicillin/Streptomycin and 10% FBS. Cells were cultured at 37 °C in a humidified 5% CO2 incubator.

### Engineered cells

All engineered cell lines were generated by lentiviral transduction. Lentiviral vectors were generated using a third-generation lentiviral packaging system (gift of D. Baltimore). Constructs encoding the lentiviral packaging plasmids and helper plasmids were transfected into HEK293T cells with PEI. 68 hours later, viral supernatant was collected from HEK293T cells and concentrated using the Lenti-X Concentrator system (Takara Bio Inc.). Concentrated virus was then applied to RAW 264.7 macrophages in the presence of RetroNectin infection reagent (Takara Bio) and cells were spun at 1000 g for 1 hour at 32 °C. After at least 7 days of growth post-infection, cells were sorted using the MACSQuant Tyto (Miltenyi Biotec) to obtain a population of cells that were BFP+, RFP+, and GFP-, ensuring that cells had both components of the circuit and removing any cells that had been activated during the lentiviral transduction process.

### Heat induction assays

Heat induction of transduced RAW 264.7 cells was performed in a BioRad C1000 Thermocycler. Cells were resuspended at 4 million/ml and then 50 µl was transferred to a sterile PCR tube for heating. Immediately after heat induction, cells were plated and returned to normal mammalian culture conditions. At each timepoint, cells were scraped and resuspended in PBS for flow cytometry. Fluorescent reporters were measured using the MACSQuant Analyzer 10 Flow Cytometer (Miltenyi Biotec). For viability and growth rate studies, cells were stained using 7-AAD and assessed 24 hours after heating. Three biological replicates were performed for all assays.

### Animal procedures

Female BALB/cJ mice aged 8-12 weeks were purchased from Jackson Laboratory. Animal experiments were approved by the Caltech Institutional Animal Care and Use Committee (protocol 1697). All researchers involved in animal experiments complied with relevant animal-use guidelines.

#### Tumor implantation

To establish the tumor model, mice were anesthetized using a 1-2% isoflurane-air mixture. The injection site was shaved and sterilized using an isopropyl wipe, and a total of 2 million cells (1:100 4T1:RAW-togLuc) were resuspended in 100 µl Matrigel and implanted subcutaneously in the flank.

#### *In vivo* ultrasound

Mice were anesthetized using a 1-2% isoflurane-air mixture and placed on 37 °C heat pad. Respiration rate was maintained at 50-100 breaths per minute throughout the ultrasound procedure. Hair removal cream (Nair) was applied to the tumor and surrounding skin to remove fur. A SoundCare Plus ultrasound device (Richmar) equipped with a 1 cm^2^ transducer was used at 3 MHz center frequency, 100% duty cycle, and 0.1-2 W/cm^2^ output power for 15 minutes. The peak-to-peak pressure at the focal point was 516 kPa given an applied power of 2 W/cm^2^, as measured using a needle hydrophone (Onda HNR-0500) in a water bath. Output power was manually adjusted to maintain tumor skin temperature at 43 °C. Skin temperature was monitored using a thermal camera (Teledyne FLIR). A layer of ultrasound gel at least 5 mm thick was applied between the transducer and the tumor to position the tumor in the focal zone of the ultrasound beam.

#### In vivo bioluminescence imaging

Bioluminescence imaging of mice was performed using an IVIS Lumina LT Series III (Revvity). For AkaLuc imaging, 100 μl 5 mM Tokeoni (Sigma-Aldrich) was administered intraperitoneally, and images were acquired 20 minutes after injection. For Antares imaging, 50 μl 8.8 mM Ffz (Promega) was administered intraperitoneally, and images were acquired 15 minutes after injection. AkaLuc and Antares imaging was performed 6-8 hours apart. Images were analyzed using Living Image software (Revvity).

#### Flow cytometry and histology of tumors

Flow cytometry: Tissues were harvested, mechanically dissociated by chopping with a razor blade, and transferred to a digestion buffer (Leibovitz’s media with 0.1 mg/mL DNase I and 2 mg/mL Collagenase P). The tissue samples were incubated at 37°C for 1 hour with continuous rotation. After incubation, samples were washed twice with flow buffer (HBSS with 0.25% BSA), filtered through a 70 µm cell strainer, stained with SYTOX Blue dead cell stain (Invitrogen), and analyzed by flow cytometry (MacsQuant Analyzer 10).

Histology: Tumors were extracted and fixed overnight in 4% paraformaldehyde at 4°C with continuous rotation. They were then immersed in 30% sucrose for 48 hours before embedding in O.C.T. Compound (Fisher Scientific) at −80°C. The frozen tissue was cryosectioned at 20 µm thickness and stained with DAPI (Thermo Scientific). Images were acquired using a STELLARIS confocal microscope (Leica).

### IL-12 quantification

All IL-12 measurements were quantified using the Mouse IL-12 p70 DuoSet ELISA kit (R&D Systems DY419) according to manufacturer’s instructions. Concentrations were calculated using a four-parameter logistic standard curve and technical duplicates were performed for all assays. To calculate the IL-12 produced per cell per day (Figure 4b), cells were induced as above. Three days after induction, 5×10^5^ cells were seeded in 1 ml media, and media was collected 24 hours later. To measure IL-12 production over 43 days (Figure 4c), cells were induced as above. 5×10^4^ cells were seeded in 1 ml of media 3 days prior to media collection at each timepoint. Media was collected and stored at −80°C until all time points were acquired. IL-12 measurements were normalized to cell count and the peak IL-12 measurement.

## ACKNOWLEDGMENTS

The authors thank Dr. Will Benman for assistance with heating equipment and Dr. Alen Pavlic for help with transducer calibration. Figures [1a, 2e, 3b, 4a and S5a] were created with BioRender.com. Confocal microscopy was performed in the Caltech Beckman Institute Biological Imaging Facility with help from Dr. Giada Spigolon and Dr. Andres Collazo. This research was supported by the Sontag Foundation, the National Institutes of Health (DP1 EB033154 to M.G.S.), the Jacobs Institute for Molecular Engineering for Medicine, and the DeepMIC center of the Beckman Institute at Caltech. A.S.F. was supported by the Natural Sciences and Engineering Research Council of Canada. M.H.A. was supported by the National Science Foundation graduate research fellowship and the Paul and Daisy Soros Fellowship for New Americans. C.A.B.S was supported by the International Human Frontier Science Program Organization (grant LT0036/2022-L). M.G.S. is an investigator of the Howard Hughes Medical Institute.

## AUTHOR CONTRIBUTIONS

A.L., A.S.F., M.H.A., and M.G.S. conceived and designed the study. A.L., A.S.F., M.H.A., C.A.B.S., and D.W. planned and conducted experiments. A.L. and A.S.F. analyzed the data. A.L., A.S.F., and M.G.S. wrote the manuscript with input from all other authors. M.G.S. supervised the research.

## NOTES

A.L., A.S.F., M.H.A. and M.G.S. are inventors on a patent application filed by Caltech pertaining to this work (PCT/US2022/077483). MGS is a co-founder of Port Therapeutics.

## SUPPLEMENTARY MATERIALS

Figures S1−S6: (i) RAW-togFP expression of OFF- and ON-state reporters over five days, (ii) heat induction effect on transgene stability, (iii) characterization of the unfocused 3 MHz ultrasound transducer, (iv) 4T1 + RAW-togLuc tumor growth curves and AkaLuc expression over time, (v) production of IL-12 from RAW-Ctrl cells after heat induction, (vi) RAW-togIL12 constitutive and OFF-state reporter expression.

## REFERENCES

(1) Cappell, K. M., and Kochenderfer, J. N. (2023) Long-term outcomes following CAR T cell therapy: what we know so far. Nat. Rev. Clin. Oncol. 20, 359–371.

(2) Neelapu, S. S., Locke, F. L., Bartlett, N. L., Lekakis, L. J., Miklos, D. B., Jacobson, C. A., Braunschweig, I., Oluwole, O. O., Siddiqi, T., Lin, Y., Timmerman, J. M., Stiff, P. J., Friedberg, J. W., Flinn, I. W., Goy, A., Hill, B. T., Smith, M. R., Deol, A., Farooq, U., McSweeney, P., Munoz, J., Avivi, I., Castro, J. E., Westin, J. R., Chavez, J. C., Ghobadi, A., Komanduri, K. V., Levy, R., Jacobsen, E. D., Witzig, T. E., Reagan, P., Bot, A., Rossi, J., Navale, L., Jiang, Y., Aycock, J., Elias, M., Chang, D., Wiezorek, J., and Go, W. Y. (2017) Axicabtagene Ciloleucel CAR T-Cell Therapy in Refractory Large B-Cell Lymphoma. N. Engl. J. Med. 377, 2531–2544.

(3) Weber, E. W., Maus, M. V., and Mackall, C. L. (2020) The Emerging Landscape of Immune Cell Therapies. Cell 181, 46–62.

(4) Schett, G., Mackensen, A., and Mougiakakos, D. (2023) CAR T-cell therapy in autoimmune diseases. The Lancet 402, 2034–2044.

(5) Mackensen, A., Müller, F., Mougiakakos, D., Böltz, S., Wilhelm, A., Aigner, M., Völkl, S., Simon, D., Kleyer, A., Munoz, L., Kretschmann, S., Kharboutli, S., Gary, R., Reimann, H., Rösler, W., Uderhardt, S., Bang, H., Herrmann, M., Ekici, A. B., Buettner, C., Habenicht, K. M., Winkler, T. H., Krönke, G., and Schett, G. (2022) Anti-CD19 CAR T cell therapy for refractory systemic lupus erythematosus. Nat. Med. 28, 2124–2132.

(6) Labani-Motlagh, A., Ashja-Mahdavi, M., and Loskog, A. (2020) The Tumor Microenvironment: A Milieu Hindering and Obstructing Antitumor Immune Responses. Front. Immunol. 11.

(7) Yeku, O. O., Purdon, T. J., Koneru, M., Spriggs, D., and Brentjens, R. J. (2017) Armored CAR T cells enhance antitumor efficacy and overcome the tumor microenvironment. Sci. Rep. 7, 10541.

(8) Li, X., Shao, M., Zeng, X., Qian, P., and Huang, H. (2021) Signaling pathways in the regulation of cytokine release syndrome in human diseases and intervention therapy. Signal Transduct. Target. Ther. 6, 1–16.

(9) Wang, V., Gauthier, M., Decot, V., Reppel, L., and Bensoussan, D. (2023) Systematic Review on CAR-T Cell Clinical Trials Up to 2022: Academic Center Input. Cancers 15, 1003.

(10) Piraner, D. I., Farhadi, A., Davis, H. C., Wu, D., Maresca, D., Szablowski, J. O., and Shapiro, M. G. (2017) Going Deeper: Biomolecular Tools for Acoustic and Magnetic Imaging and Control of Cellular Function. Biochemistry 56, 5202–5209.

(11) Tan, P., He, L., Han, G., and Zhou, Y. (2017) Optogenetic Immunomodulation: Shedding Light on Antitumor Immunity. Trends Biotechnol. 35, 215–226.

(12) Abedi, M. H., Lee, J., Piraner, D. I., and Shapiro, M. G. (2020) Thermal Control of Engineered T-cells. ACS Synth. Biol. 9, 1941–1950.

(13) Miller, I. C., Zamat, A., Sun, L.-K., Phuengkham, H., Harris, A. M., Gamboa, L., Yang, J., Murad, J. P., Priceman, S. J., and Kwong, G. A. (2021) Enhanced intratumoural activity of CAR T cells engineered to produce immunomodulators under photothermal control. Nat. Biomed. Eng. 5, 1348–1359.

(14) Wu, Y., Liu, Y., Huang, Z., Wang, X., Jin, Z., Li, J., Limsakul, P., Zhu, L., Allen, M., Pan, Y., Bussell, R., Jacobson, A., Liu, T., Chien, S., and Wang, Y. (2021) Control of the activity of CAR-T cells within tumours via focused ultrasound. Nat. Biomed. Eng. 5, 1336–1347.

(15) ter Haar, G., and Coussios, C. (2007) High intensity focused ultrasound: physical principles and devices. Int. J. Hyperth. Off. J. Eur. Soc. Hyperthermic Oncol. North Am. Hyperth. Group 23, 89–104.

(16) Abedi, M. H., Yao, M. S., Mittelstein, D. R., Bar-Zion, A., Swift, M. B., Lee-Gosselin, A., Barturen-Larrea, P., Buss, M. T., and Shapiro, M. G. (2022) Ultrasound-controllable engineered bacteria for cancer immunotherapy. Nat. Commun. 13, 1585.

(17) Pittet, M. J., Michielin, O., and Migliorini, D. (2022) Clinical relevance of tumour-associated macrophages. Nat. Rev. Clin. Oncol. 19, 402–421.

(18) Klichinsky, M., Ruella, M., Shestova, O., Lu, X. M., Best, A., Zeeman, M., Schmierer, M., Gabrusiewicz, K., Anderson, N. R., Petty, N. E., Cummins, K. D., Shen, F., Shan, X., Veliz, K., Blouch, K., Yashiro-Ohtani, Y., Kenderian, S. S., Kim, M. Y., O’Connor, R. S., Wallace, S. R., Kozlowski, M. S., Marchione, D. M., Shestov, M., Garcia, B. A., June, C. H., and Gill, S. (2020) Human chimeric antigen receptor macrophages for cancer immunotherapy. Nat. Biotechnol. 38, 947–953.

(19) Brempelis, K. J., Cowan, C. M., Kreuser, S. A., Labadie, K. P., Prieskorn, B. M., Lieberman, N. A. P., Ene, C. I., Moyes, K. W., Chinn, H., DeGolier, K. R., Matsumoto, L. R., Daniel, S. K., Yokoyama, J. K., Davis, A. D., Hoglund, V. J., Smythe, K. S., Balcaitis, S. D., Jensen, M. C., Ellenbogen, R. G., Campbell, J. S., Pierce, R. H., Holland, E. C., Pillarisetty, V. G., and Crane, C. A. (2020) Genetically engineered macrophages persist in solid tumors and locally deliver therapeutic proteins to activate immune responses. J. Immunother. Cancer 8, e001356.

(20) Xue, Y., Yan, X., Li, D., Dong, S., and Ping, Y. (2024) Proinflammatory polarization of engineered heat-inducible macrophages reprogram the tumor immune microenvironment during cancer immunotherapy. Nat. Commun. 15, 2270.

(21) Deckers, R., Quesson, B., Arsaut, J., Eimer, S., Couillaud, F., and Moonen, C. T. W. (2009) Image-guided, noninvasive, spatiotemporal control of gene expression. Proc. Natl. Acad. Sci. U. S. A. 106, 1175–1180.

(22) Schnütgen, F., Doerflinger, N., Calléja, C., Wendling, O., Chambon, P., and Ghyselinck, N. B. (2003) A directional strategy for monitoring Cre-mediated recombination at the cellular level in the mouse. Nat. Biotechnol. 21, 562–565.

(23) Kay, R. J., Boissy, R. J., Russnak, R. H., and Candido, E. P. (1986) Efficient transcription of a Caenorhabditis elegans heat shock gene pair in mouse fibroblasts is dependent on multiple promoter elements which can function bidirectionally. Mol. Cell. Biol. 6, 3134–3143.

(24) Evans, S. S., Repasky, E. A., and Fisher, D. T. (2015) Fever and the thermal regulation of immunity: the immune system feels the heat. Nat. Rev. Immunol. 15, 335–349.

(25) Corish, P., and Tyler-Smith, C. (1999) Attenuation of green fluorescent protein half-life in mammalian cells. Protein Eng. 12, 1035–1040.

(26) Hildebrandt, B., Wust, P., Ahlers, O., Dieing, A., Sreenivasa, G., Kerner, T., Felix, R., and Riess, H. (2002) The cellular and molecular basis of hyperthermia. Crit. Rev. Oncol. Hematol. 43, 33–56.

(27) Chu, K. F., and Dupuy, D. E. (2014) Thermal ablation of tumours: biological mechanisms and advances in therapy. Nat. Rev. Cancer 14, 199–208.

(28) Chu, J., Oh, Y., Sens, A., Ataie, N., Dana, H., Macklin, J. J., Laviv, T., Welf, E. S., Dean, K. M., Zhang, F., Kim, B. B., Tang, C. T., Hu, M., Baird, M. A., Davidson, M. W., Kay, M. A., Fiolka, R., Yasuda, R., Kim, D. S., Ng, H.-L., and Lin, M. Z. (2016) A bright cyan-excitable orange fluorescent protein facilitates dual-emission microscopy and enhances bioluminescence imaging in vivo. Nat. Biotechnol. 34, 760–767.

(29) Iwano, S., Sugiyama, M., Hama, H., Watakabe, A., Hasegawa, N., Kuchimaru, T., Tanaka, K. Z., Takahashi, M., Ishida, Y., Hata, J., Shimozono, S., Namiki, K., Fukano, T., Kiyama, M., Okano, H., Kizaka-Kondoh, S., McHugh, T. J., Yamamori, T., Hioki, H., Maki, S., and Miyawaki, A. (2018) Single-cell bioluminescence imaging of deep tissue in freely moving animals. Science 359, 935–939.

(30) Su, Y., Walker, J. R., Park, Y., Smith, T. P., Liu, L. X., Hall, M. P., Labanieh, L., Hurst, R., Wang, D. C., Encell, L. P., Kim, N., Zhang, F., Kay, M. A., Casey, K. M., Majzner, R. G., Cochran, J. R., Mackall, C. L., Kirkland, T. A., and Lin, M. Z. (2020) Novel NanoLuc substrates enable bright two-population bioluminescence imaging in animals. Nat. Methods 17, 852–860.

(31) Nguyen, K. G., Vrabel, M. R., Mantooth, S. M., Hopkins, J. J., Wagner, E. S., Gabaldon, T. A., and Zaharoff, D. A. (2020) Localized Interleukin-12 for Cancer Immunotherapy. Front. Immunol. 11.

(32) Cabrera, A., Edelstein, H. I., Glykofrydis, F., Love, K. S., Palacios, S., Tycko, J., Zhang, M., Lensch, S., Shields, C. E., Livingston, M., Weiss, R., Zhao, H., Haynes, K. A., Morsut, L., Chen, Y. Y., Khalil, A. S., Wong, W. W., Collins, J. J., Rosser, S. J., Polizzi, K., Elowitz, M. B., Fussenegger, M., Hilton, I. B., Leonard, J. N., Bintu, L., Galloway, K. E., and Deans, T. L. (2022) The sound of silence: Transgene silencing in mammalian cell engineering. Cell Syst. 13, 950–973.

(33) Di Stasi, A., Tey, S.-K., Dotti, G., Fujita, Y., Kennedy, Nasser Alana, Martinez, C., Straathof, K., Liu, E., Durett, A. G., Grilley, B., Liu, H., Cruz, C. R., Savoldo, B., Gee, A. P., Schindler, J., Krance, R. A., Heslop, H. E., Spencer, D. M., Rooney, C. M., and Brenner, M. K. (2011) Inducible Apoptosis as a Safety Switch for Adoptive Cell Therapy. N. Engl. J. Med. 365, 1673–1683.

