## Supplementary Material for "Thermally controlled state switches for macrophage immunotherapy"

### SUPPLEMENTARY MATERIALS

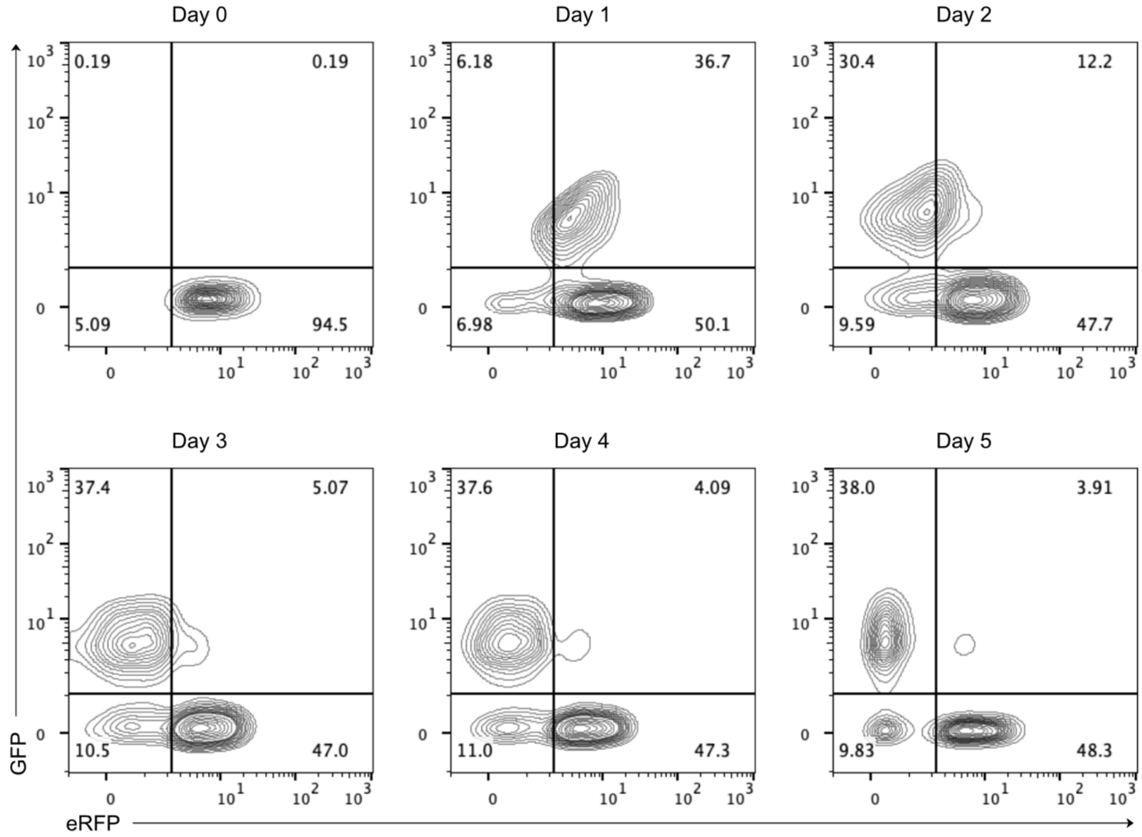


**Figure S1. RAW-togFP expression of OFF- and ON-state reporters over five days.** Activated RAW-togFP cells are double-positive (GFP+/eRFP+) one day after heat induction, and gradually shift to GFP+/eRFP− over five days. Cells were first gated for doublet discrimination and BFP+.

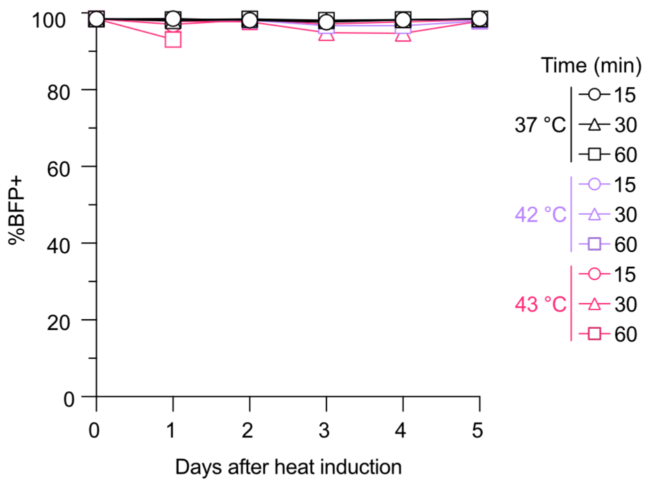


**Figure S2. Heat induction does not affect transgene stability.** Percentage of BFP+ cells, indicating presence of HSP16-Cre construct, remains stable over 5 days following heat induction.

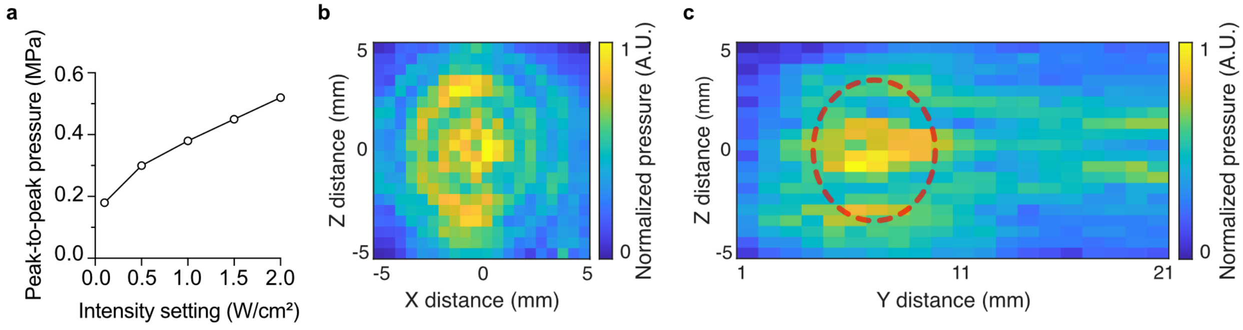


**Figure S3. Characterization of the unfocused 3 MHz ultrasound transducer.** (**a**) Calibration plot of peak-to-peak pressure at a single point in the center of the focal region of the transducer under varying applied intensity settings, measured using a needle hydrophone. (**b,c**) Normalized pressure field of the 3 MHz transducer in the transverse plane, orthogonal to direction of propagation (b), and in the longitudinal plane, along the direction of propagation (c). Red circle represents where the targeted tumor would lie within the longitudinal plane.

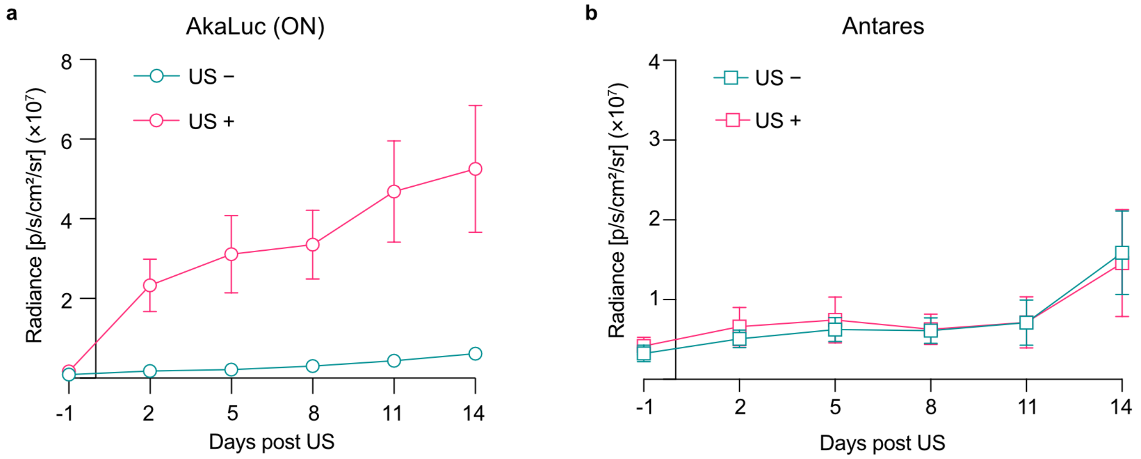


**Figure S4. 4T1 + RAW-togLuc tumor growth curves and AkaLuc expression over time.** (**a**) Quantification of AkaLuc (ON) luminescence from tumors treated with US (pink) or untreated on the opposite flank (blue). (**b**) Growth curves of 4T1+RAW-togLuc tumors, measured via quantification of Antares luminescence, treated with US (pink) or untreated on the opposite flank (blue). N = 7 mice. Bars = SEM.

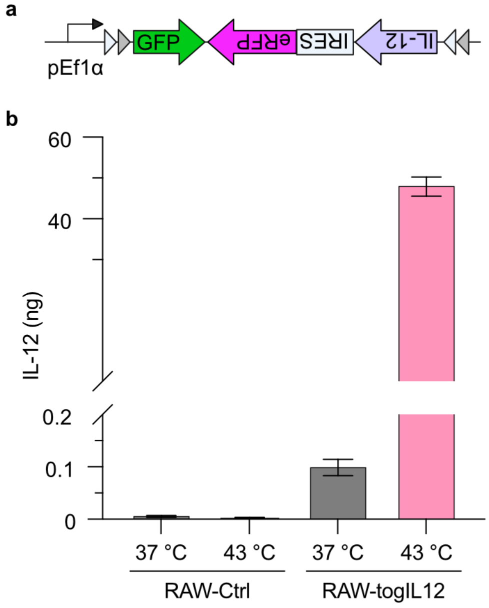


**Figure S5. RAW-Ctrl cells do not produce significant amounts of IL-12 after heat induction.** (**a**) RAW cells were transduced with only the toggle switch element of the togIL12 thermal circuit to generate RAW-Ctrl cells. The HSP16-Cre actuator element was not transduced. (**b**) IL-12 production quantified from culture media of RAW-Ctrl and RAW-togIL12 cells three days after incubation at 37 or 43 °C for 15 minutes. n = 3 replicates. Bars = SEM.

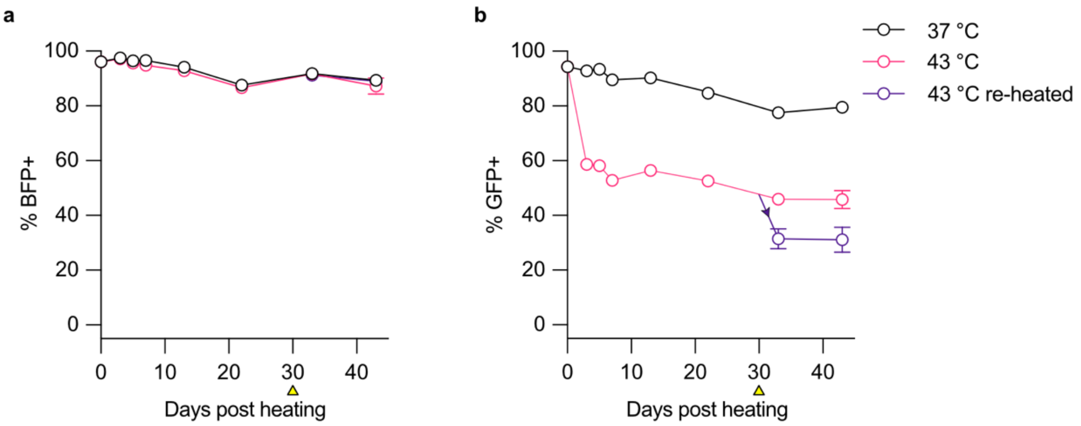


**Figure S6. RAW-togIL12 constitutive and OFF-state reporter expression.** RAW-togIL12 cells were incubated at 37 and 43 °C for 15 minutes and then analyzed via flow cytometry for 43 days. Percent BFP+ cells (**a**) and percent GFP+ of BFP+ cells (**b**) were determined after first gating for doublet discrimination. At t = 30 days (yellow arrowhead), a subpopulation of cells that had been heated at 43 °C at t = 0 were incubated again at 43 °C for 15 minutes and then further assessed on days 33 and 43 (purple lines).
